# Natural selection of synthetic gene drives for population suppression can favour an intermediate strength of drive

**DOI:** 10.1101/2024.11.03.621714

**Authors:** PJ Beaghton, Austin Burt

## Abstract

Synthetic gene drives are being investigated as tools to suppress pest populations, and it is important to understand how natural selection will act on variant drivers that may either arise by *de novo* mutation or are intentionally released. In this study we extend previous spatially implicit stochastic models to examine the evolutionary dynamics of synthetic driving Y chromosomes in patchy environments when population size is responding dynamically to the spread of the driver, and derive conditions for the existence of an evolutionarily stable strategy (ESS) for drive strength. Under broad conditions an intermediate drive strength emerges as the ESS, capable of outcompeting both stronger and weaker variants. Additionally, we show how the intentional release of two drivers straddling the ESS can help stabilise population dynamics. Finally, inbreeding depression has the effect of expanding the range of conditions under which no intermediate ESS exists, with ever stronger drive being selected until the population is eliminated. These results provide insights into the expected evolutionary trajectories of gene drive systems, with important implications for the design and release of gene drives for pest and vector control.

## Introduction

Synthetic gene drives are being investigated as potential new tools to modify or suppress disease vectors, harmful invasive species, and other pest species (Hay *et al*., 2021; Bier, 2021; Nolan, 2021). The basic idea for suppressing a target population is to insert a gene into the genome that is inherited at a greater-than-Mendelian rate and will both spread through a population and disrupt reproduction by distorting the sex ratio or causing homozygous lethality or sterility (Burt, 2003; Kyrou *et al*., 2018; Simoni *et al*., 2020). In principle, only very small releases are necessary, and the gene drive can then increase in frequency of its own accord over successive generations, gradually suppressing the population as it does so. Naturally-occurring dispersal by the target organism can extend the suppression to new locations, and in this way it can be extremely efficient, with small releases leading to large impacts (North *et al*., 2019, 2020). This efficiency can be particularly important when the target population is widespread and there are few resources to deal with it (Burt *et al*., 2018).

In simple deterministic models of this process the population is suppressed by the presence of the driver, and if the drive is strong enough the population can be eliminated (Deredec *et al*., 2008, 2011; Beaghton *et al*., 2016). However, more detailed modelling has also shown that under some circumstances, particularly when there are both stochastic and spatial dynamics, a population can persist indefinitely after release of a suppression gene drive, regardless of how strong the drive (Eckhoff *et al*., 2017; North *et al*., 2019; Champer *et al*., 2021). One scenario that allows indefinite persistence includes a combination of spatial structure, a life history in which inbreeding becomes more common at low densities, and low inbreeding depression so that the inbred progeny produced at low densities can contribute to the population (Bull *et al*., 2019; Beaghton & Burt, 2022). Persistence in these circumstances was demonstrated for both driving Y chromosomes and for autosomal drivers causing homozygous lethality.

Under those circumstances where there can be long-term persistence of a driver and the population, it is interesting to enquire how selection might act on constructs with different rates of drive. More specifically, if one releases more than one type of drive, which one(s) are expected to persist and which disappear? What will happen to mutations that arise in a construct after release and affect the rate of drive? In deterministic or non-spatial models we generally expect stronger drivers to displace weaker ones, but it is not clear that this will be the case under the circumstances described above, where the population can persist regardless of the strength of the drive. In these models the driver is selected against when population densities are low, and this effect may be expected to be stronger when the drive is stronger. Indeed, the life history that Beaghton & Burt (2022) found gives population persistence is the same as used in the local mate competition model of Hamilton (1967), in which, in the context of stable population size (i.e., population size is treated as an extrinsic parameter), a driving Y evolves to a single intermediate ‘unbeatable’ sex ratio. Here we investigate whether the same is true when population size is instead an intrinsic variable, responding dynamically to the presence of the suppressive driver. We also investigate the fate of combinations of drivers released into a population, and explore the effect of incorporating inbreeding depression on the evolutionary dynamics.

### The Model

The model is based on scenario 5 in Beaghton & Burt (2022). In brief, males are XY and either have a wildtype Y chromosome that is transmitted to 50% of sperm (and offspring) or a driving Y that is transmitted to a proportion *m* of offspring. There are no fitness differences between Y chromosomes other than through their effect on the sex ratio. The environment is divided into an infinite number of patches; individuals are born in patches; and females mate once with a randomly chosen male from the same patch, store the sperm, and disperse to another patch, randomly chosen, where they reproduce and die (Fig. 1). Generations are therefore discrete and the number of mated females settling in a patch is Poisson-distributed. Each mated female produces a Poisson-distributed number of offspring with mean 2*R/*(1 + *αF*), where *F* is the number of mated females reproducing in the patch; *a >* 0 measures the strength of density-dependence on recruitment rate; and *R/*(1 − *α*) is the maximum low-density rate of increase for the wildtype population (*m* = 0.5). The sex ratio of the progeny produced by each female is binomially-distributed with probability dependent on the Y of the male she mated with.

**Figure 1:**
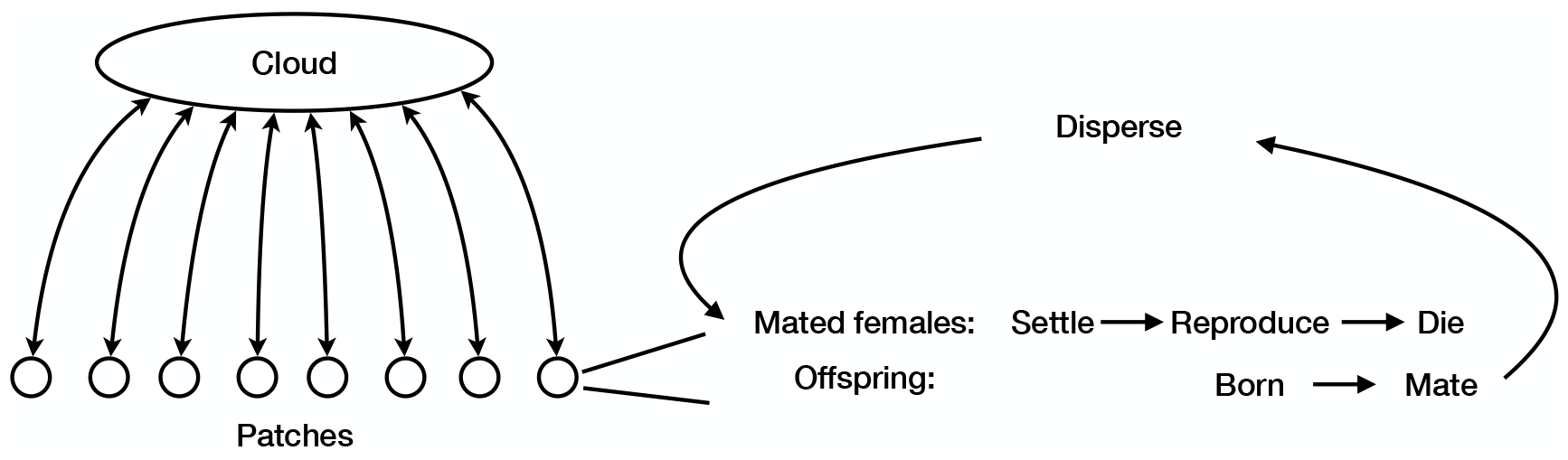
The model considers an infinite number of patches, within which a finite number of mated females settle, reproduce, and die. Their offspring mate randomly, and then mated females disperse via a well-mixed cloud to another randomly chosen patch to start the cycle again.

Here we extend this model to consider not a wild-type and a driving Y, but instead two different driving Ys with arbitrary strengths of drive, *m*_*A*_ and *m*_*B*_, where 0 ≤ *m*_*A*_, *m*_*B*_ ≤ 1. (In Appendix A we present a recurrence equations for an arbitrary number of different drives, but believe that the system will typically evolve to have at most 2 types of Y at equilibrium, and so focus on that case here.)

The recurrence equations for the population densities 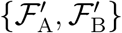 of genotypes A and B in the next generation, in terms of the population densities {ℱ_A_, ℱ_B_} in the previous generation, their respective strengths of drive *m*_A_, and *m*_B_ and the inputs *R* and *α*, are:

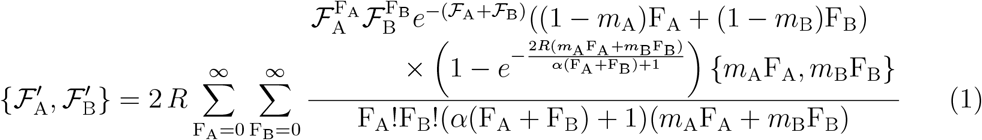

As *t* → ∞ and for given {*R, α, m*_A_, *m*_B_}, Eqs(1) may admit one or more stable and/or unstable equilibrium solutions 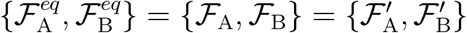 in addition to the trivial solution {0, 0}.

## The Evolutionary Stable Strategy

### Monomorphic equilibria

We postulate that for a range of *R* and *α* there exists a unique value *m*^∗^ for which genotype A with strength of drive *m*_A_ = *m*^∗^ will **always** replace genotype B with strength of drive *m*_B_ ≠ *m*^∗^ and as *t* → ∞ its population density will converge to a **stable** fixed point with equilibrium population density 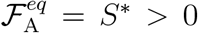. We also postulate that if neither *m*_A_ nor *m*_B_ are equal to *m*^∗^, then the following apply: (1) If both strengths of drive are below or above *m*^∗^, then at equilibrium the genotype with strength of drive closest to *m*^∗^ will be fixed (although if too strong it may still eliminate the population), (2) If the two strengths of drive straddle *m*^∗^, at equilibrium one may be fixed or both may coexist. We first derive the equations for *m*^∗^ and *S*^∗^ and then we analyze the stability of the solutions to these equations.

#### Derivation

Conditional on the postulates above being true, Eqs(1) are used to evaluate the population densities 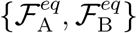 at *t* → ∞ by setting 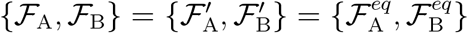 and introducing the change of variables 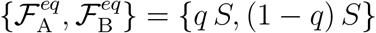 and {*m*_A_, *m*_B_} = { *m, m* + *ϵ* }, where *q* is the equilibrium fraction of genotype A, *S* is the aggregate population density of the two genotypes, *m*_A_ = *m* and *m*_B_ = *m* + *ϵ*:

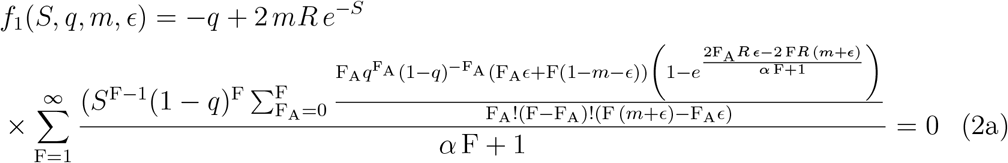

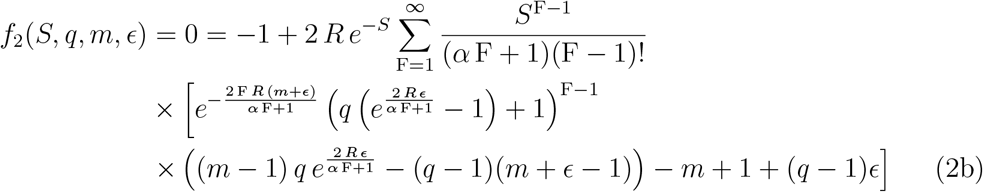

We shall now use Eqs(2a-b) to derive the equations for *m* = *m*^∗^ and the corresponding equilibrium population *S* = *S*^∗^. We start by examining the total differentials of *f*_1_(*S, q, m, ϵ*) and *f*_2_(*S, q, m, ϵ*) in Eqs (2a-b). Since the values of *f*_1_ and *f*_2_ remain unchanged and equal to 0 for all acceptable solutions *S, q, m, ϵ*, it follows that all the solutions of Eqs(2a-b) satisfy *df*_1_ = *df*_2_ = 0:

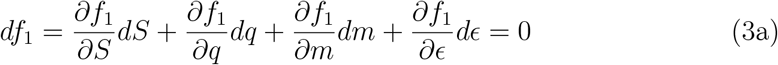

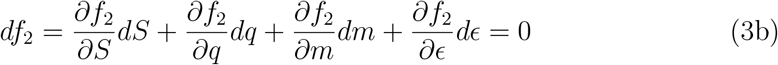

We solve the equations above for *dq*:

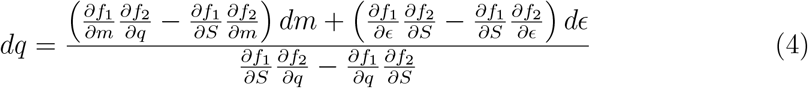

We are looking for values of *m* that are fixed at equilibrium (i.e. *q* = 1) regardless of whether the other allele has a rate of drive *m* + *ϵ* higher or lower than *m* (i.e. whether *ϵ* is positive or negative). Furthermore, when *q* = 1, for any value of *m* ≠m^∗^, a change of *ϵ* from infinitesimally below (above) 0 to infinitesimally above (below) 0 changes the value *q* of the equilibrium fraction of genotype A. It is only when *m* = *m*^∗^ that a change of *ϵ* from (to) 0-to (from) 0+ has no impact on *q* (i.e. *dq* = 0 and *q* = 1) since genotype A would fix in either case. In order for this to hold, the coefficient of *dϵ* in Eq(4) must be equal to zero for the specific case *q* = 1, *ϵ* = 0 (and *m* = *m*^∗^):

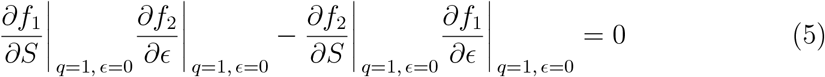

(we will henceforth mean *m*^∗^ every time we use *m*).

We now incorporate the definitions of *f*_1_ and *f*_2_ from Eqs(2a-b) into Eq(5) and after several steps in Wolfram Mathematica (including a step that shows that 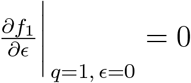), we obtain two solutions:

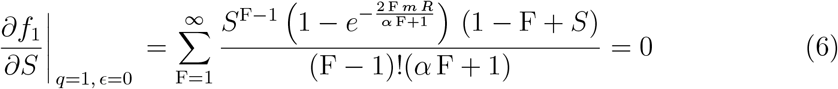

and

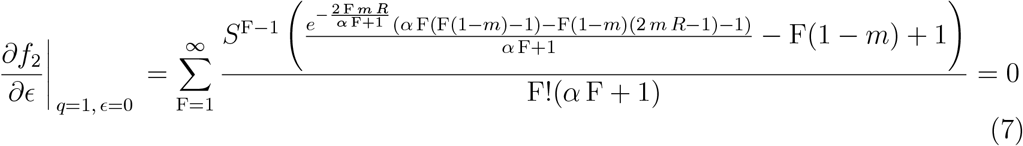

Eq(6) is only satisfied by *S* = 0 and can be disregarded.

As per our postulates, when *m* = *m*^∗^, genotype A fixes and *q* = 1. Eq(2b) then becomes

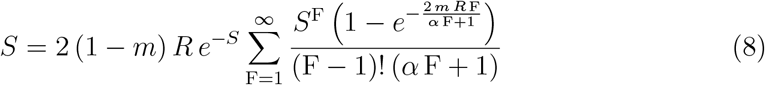

This equation gives the equilibrium population size(s) as a function of *m* when there is only a single type of Y chromosome in the population. Depending on the values of *R* and *α*, for any given *m* the single genotype equation admits (beyond the trivial solution *S* = 0) (i) no solution for *S >* 0, (ii) two non-zero solutions for *S*, one unstable, representing an Allee critical threshold, and another, higher and stable, or (iii) one stable non-zero solution. The stability analysis of the single genotype system is presented in Appendix B.

#### Stability analysis

Having identified a unique pair {*m*^∗^, *S*^∗^} for any given *R* and *α*, we now examine the linear stability of the equilibrium solution 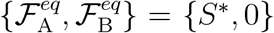, where A and B are genotypes with *m*_A_ = *m*^∗^ and *m*_B_ = *m*^∗^ + *ϵ*, respectively. This will allow us to ascertain under what conditions the equilibrium solution {*S*^∗^, 0} with *m* = *m*^∗^ is linearly stable to perturbations either to 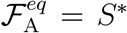 and/or to 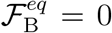. To do this, we derive the Jacobian matrix of the RHS of Eqs(2a-2b) wrt ℱ_A_ and ℱ_B_ and evaluate the matrix at the fixed point 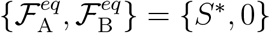. We thus obtain two real eigenvalues:

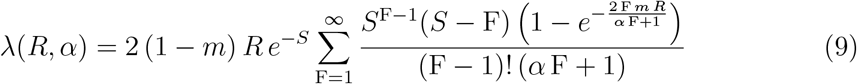

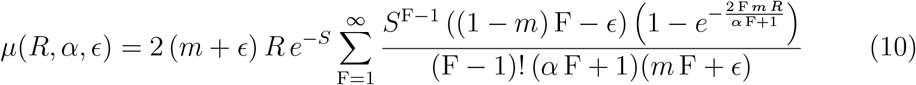

with both *m* = *m*^∗^ and *S* = *S*^∗^ also being functions of *R* and *α*.

#### Results

Eqs(7-8) are solved numerically for *m*(= *m*^∗^) and the corresponding equilibrium population *S* = *S*^∗^ for given values of the inputs *R* and *α*. These numerical solutions show that *m* = *m*^∗^ increases with *R* and decreases with *α* (Fig. 2a). The equilibrium population size also increases with *R* and decreases with *α* (Fig. 2b), but (due to the increases in *m*), it changes much less than the pre-release equilibrium population size (Fig. 2c), and so the extent of suppression is higher for high *R* and low *α* (Fig. 2d). We can also investigate the relationship between *m* and *S* across the different values of *R* and *α*, and it turns out to be well described by Hamilton’s (1967) result of *m* = (*S* − 1)*/S*, with a maximum difference of 2% (as long as *R* ≥ 2).

**Figure 2:**
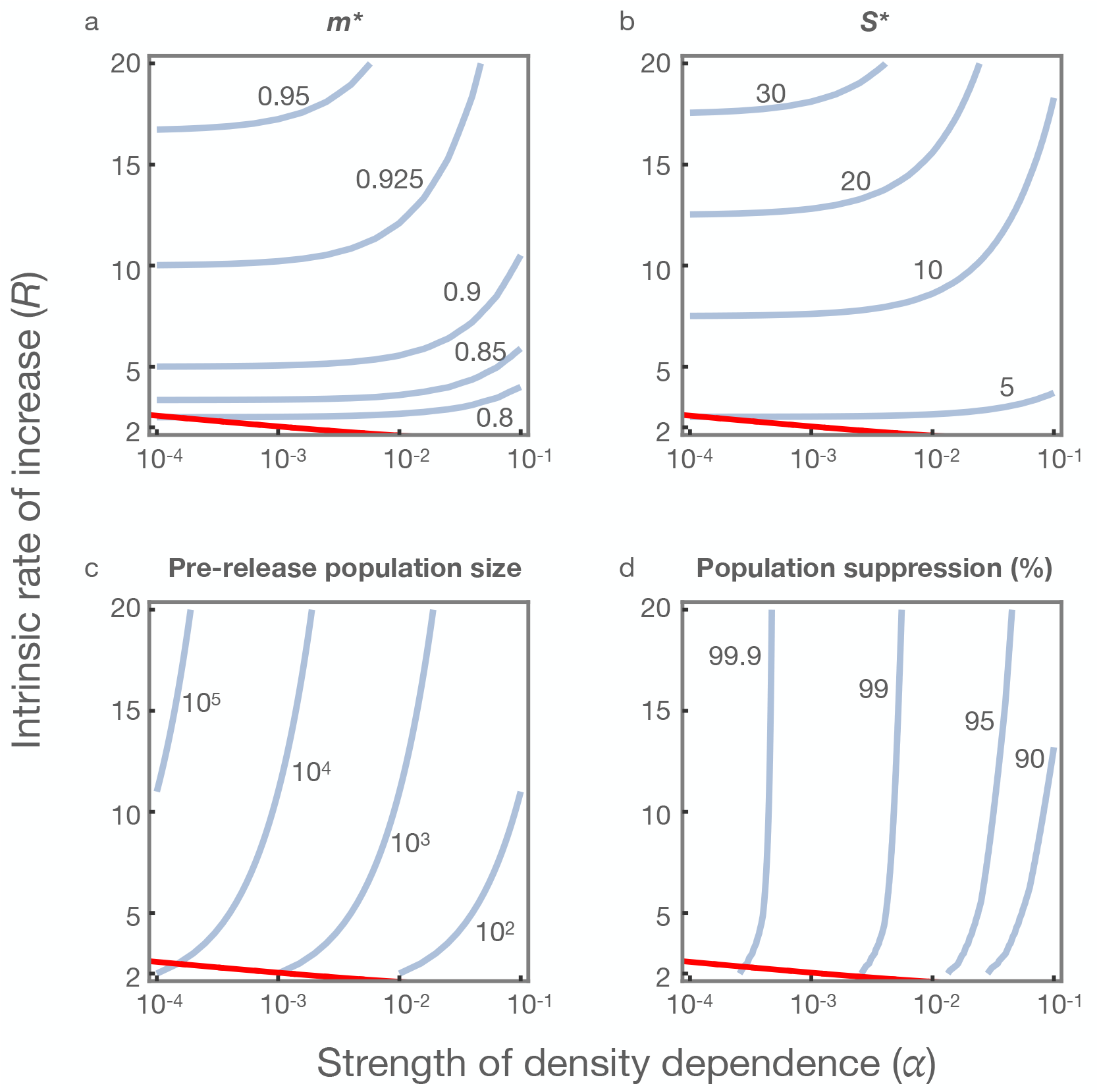
Contour plots for the equilibrium sex ratio (a) and population size (b) as a function of the intrinsic rate of increase (*R*) and the strength of density dependence (*α*). Also shown are contour plots for the pre-release population size (c) and the equilibrium extent of population suppression (d). The red lines separate the regions where the equilibrium {*m*^∗^, *S*^∗^} pair is stable (above the line) vs unstable (below).

In terms of stability, it can be easily shown that the magnitude of the eigenvalue *μ*(*R, α, ϵ*), directly related to genotype B, tends to 1 from below as *ϵ* → 0. We have also calculated *μ*(*R, α, ϵ*) numerically for a wide range of *R* and *α* and found that | *μ*(*R, α, ϵ*) | ≤ 1 for all admissible values of *ϵ*, i.e. − *m* ≤*ϵ* ≤1 − *m*. This means that an addition of a small amount of genotype B with *m*_B_ = *m*^∗^ + *ϵ* does not destabilise the equilibrium, i.e. the B perturbation dies out.

On the other hand, the magnitude of the other eigenvalue *λ*(*R, α*), directly related to the genotype A, can be greater or less than 1, depending on the values of *R* and *α*. When | *λ*| *>* 1, a small increase (or decrease) in the amount of genotype A destabilises the equilibrium and the population density of A goes to a higher stable equilibrium value (or zero). The red lines in Figure 2a-d represents the boundary between the stable and unstable regions (above and below the lines, respectively). Over most of the biologically relevant parameter space the {*m*^∗^, *S*^∗^} pair is stable, and *m*^∗^ represents an evolutionarily stable strategy (ESS), but there is a smaller region where it is unstable. In the latter region, at *m* = *m*^∗^ the equilibrium population size *S*^∗^ is unstable – that is, there is an Allee effect (Courchamp *et al*., 2008) and the population either goes to 0 (if it starts below *S*^∗^) or it goes to another (higher) equilibrium value (if it starts above *S*^∗^). This higher and stable equilibrium does NOT satisfy Eq(7).

To further investigate the differences between these two regions we calculated the selection gradient for *m*, assuming the population size *S* is at the stable equilibrium in the single genotype model (provided a solution *S >* 0 of Eq(8) exists). The selection gradient for *m* is defined as the slope of the selection coefficient on *m*, and is calculated from changes in frequency over a single generation (using Eqs(22) in the Appendix C) as

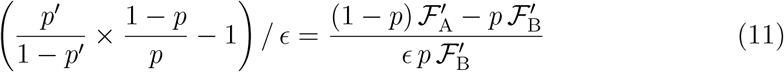

where *p* is the frequency of the stronger driver A in one generation, *p*^′^ its frequency in the next generation, and *ϵ* is the difference in rate of drive. More specifically, we set {*m*_*A*_, *m*_*B*_} = {*m* + *ϵ/*2, *m* − *ϵ/*2}, *ϵ* = 10^−6^ and {ℱ_A_, *ℱ*_B_ }= {*S p, S* (1− *p*) }, where *S* is the stable equilibrium population size associated with the value of *m* (from Eq(8)). A positive selection gradient indicates selection favours increased *m*, whereas a negative value indicates selection is for smaller *m*, assuming differences in *m* are small.

We first performed this analysis for parameter values in the stable region (*R* = 6, *α* = 0.01), and, as expected, the selection gradient is positive over much of the range, indicating selection for higher values of *m*, and then it goes negative precisely when *m* equals the previously calculated *m*^∗^ (Fig. 3a). This equality is proven analytically in Appendix C. As a further check, we simulated a population initially at {*m*^∗^, *S*^∗^}, introduced a small fraction (0.1%) of Y chromosomes with a slightly higher value of *m*, and tracked the dynamics over time (Fig. 3b). As expected, the introduced variant was selected against, decreasing monotonically in frequency, and population size remained largely unchanged.

**Figure 3:**
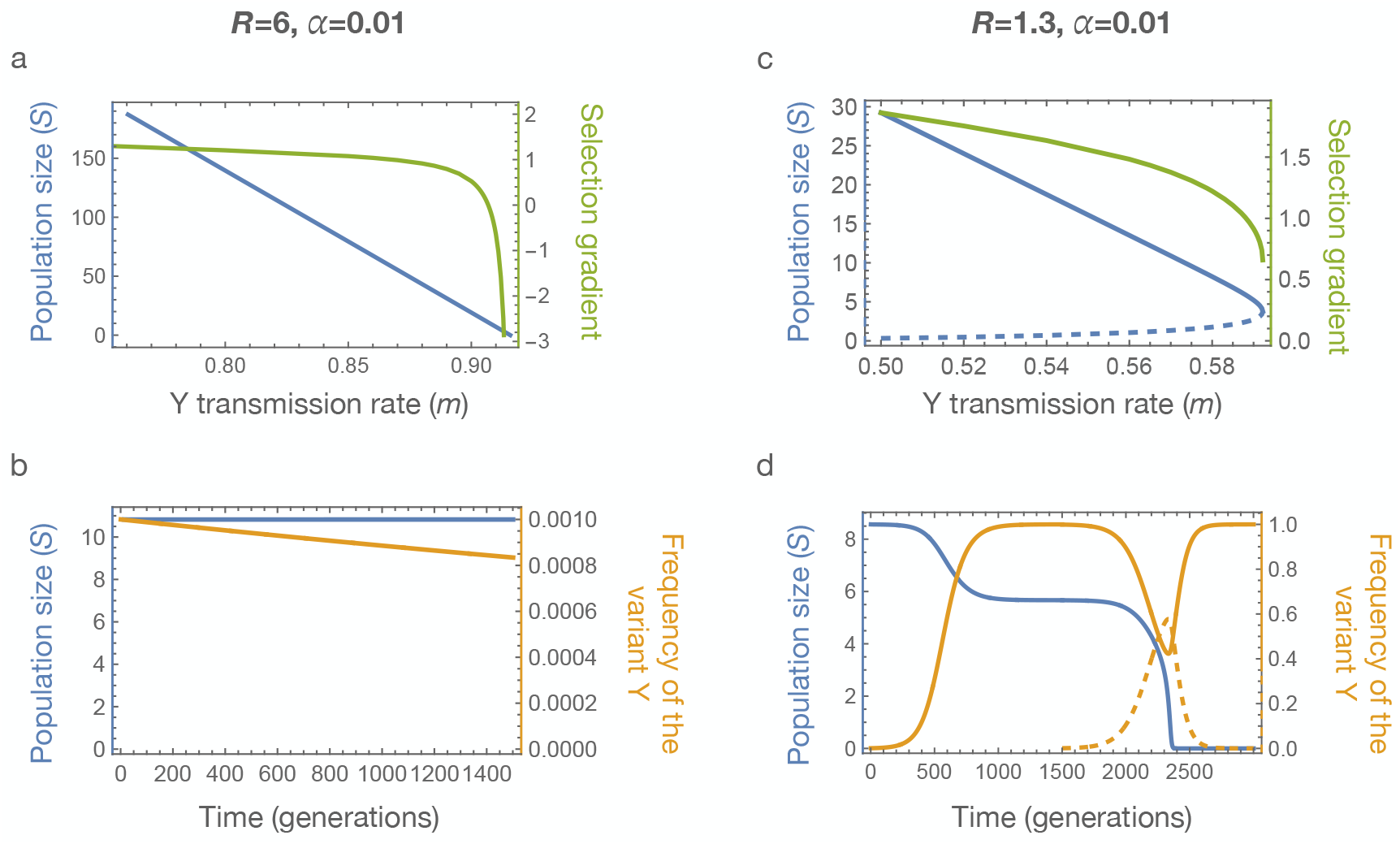
Selection gradient and dynamic analysis of stable and unstable equilibria. (a) Blue line shows the equilibrium population size as a function of the Y chromosome transmission rate (assuming it is fixed in the population), and the green line shows the selection gradient in favour of stronger drive, which declines with increasing transmission rate *m* because of the latter’s effect on population size. Note the selection gradient turns negative when *m* = *m*^∗^ as proven in Appendix C. (b) Time series showing population size (blue line) and the frequency of the stronger driver (orange line) over time. Populations start at the ESS values of *m* and *S*, and then 0.1% of males are replaced with males carrying a Y with a higher rate of drive (difference=0.01). The frequency declines over time, and the population size remains approximately constant. (c) and (d) show equivalent plots for *R* = 1.3, *α* = 0.01. Note in (c) that there are two equilibrium population sizes for each value of *m*, a higher stable equilibrium (solid line) and a lower unstable equilibrium (dashed line). The population cannot persist for values of *m* higher than the value at which the lines meet, and the selection gradient remains positive throughout this range when the population is at the biologically realistic higher equilibrium. The maximum value of *m* (and the corresponding *S >* 0) for which population is not zero are the solutions of Eqs(20-21) in Appendix B3. Note in (d) that a Y with a rate of drive 0.01 larger than *m*^∗^ increases in frequency and by generation 1500 is close to fixation, with a consequent reduction in population size; at this point another variant is introduced with drive 0.01 greater than the first variant, and it too increases in frequency (dashed line), leading to a further reduction in population size. At some point selection for the second variant is reversed, and its frequency decreases, but by then the population size is below the unstable equilibrium and it declines to zero.

We then repeated these analyses for parameter values in the unstable region (*R* = 1.3, *α* = 0.01). In this case the selection gradient (evaluated for different values of *m* and the corresponding **stable** population size, where we expect populations to be found) remains positive right up to the point where *m* is large enough to eliminate the population, never falling below 0 (Fig. 3c), indicating that there will be selection for ever higher values of *m* until the population is eliminated. To confirm this interpretation, we again simulated a population with *m* = *m*^∗^, but the population size at the higher stable equilibrium value, and again introduced a small proportion of Ys with a slightly higher *m*, and tracked the dynamics over time (Fig. 3d). In this case the variant Y with the higher *m* did not slowly disappear, but instead increased in frequency, and went to fixation, resulting in a slightly smaller population size. We then introduced another variant Y, again with a slightly higher *m*, and it too initially increased in frequency, leading to a further reduction in population size. At some point, once the population size was small enough, selection reversed, favouring the Y with the weaker drive, but by this time the population size was below the unstable equilibrium value (the Allee threshold), and it continued to decline to zero.

## Ecologically stable polymorphisms

The analysis above indicates that, for most values of *R* and *α*, and given an appropriate source of mutational variation (and in the absence of evolution by the rest of the genome), a driving Y will be expected to eventually evolve to an ESS rate of transmission. However, this could take a long time (depending on how close a released Y is to the ESS and on mutation rates), and it is therefore also interesting to know how the population will evolve in the short term, if two (or more) drivers are released that differ in their rate of transmission. If mutation rates are low, the system should relatively rapidly go to an equilibrium mixture or combination of the different types (sometimes called an evolutionarily stable state; Maynard Smith, 1982), before the slower evolution of the monomorphic ESS by selection of *de novo* mutations. The case of particular interest is where the two values of *m* straddle the ESS value, in which case we expect the wild-type Y to disappear (because it is further from the ESS than the lower of the two released drivers), and we focus on that case here.

For specific values of *R* and *α* (6 and 0.01, respectively), Figure 4a shows for different pairs of drivers, one above the ESS and one below, the frequency of the higher driver at equilibrium. As might be expected, when the two drivers are about equally different from the ESS, then they are about equal in frequency, and otherwise whichever one is closer to the ESS is more frequent.

**Figure 4:**
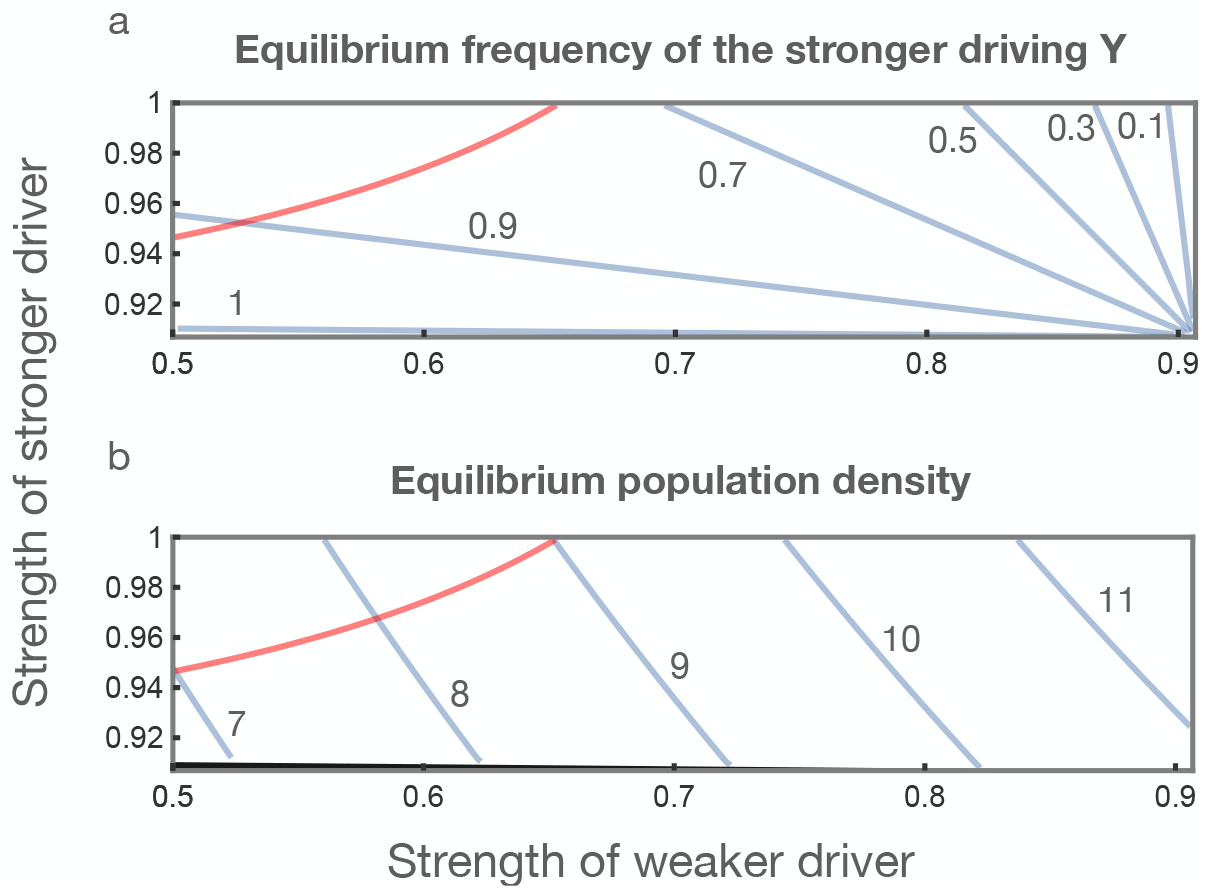
The effect of varying the strength of drive when there are two drivers straddling the ESS. Contour plots show (a) the equilibrium frequency of the stronger driver and (b) the equilibrium population size as a function of the strengths of the two drivers. Calculations are for *R* = 6 and *α* = 0.01, for which *m*^∗^ is 0.907 and *S*^∗^ is 10.8. The red line in each plot shows the Neimark-Sacker bifurcation points separating regions where the population tends to a stable equilibrium (below the curve) vs. a limit cycle (above), and the thin gray wedge along the bottom of (b) indicates the region where the stronger driver goes to fixation.

For the particular combination of parameters chosen, the equilibrium population density varies little, from about 6.9 to 11.7 mated females per deme, and in all cases is much smaller than the pre-release equilibrium (≈ (*R* − 1)*/α* = 500; Fig. 4b). When there are two drivers straddling the ESS, the equilibrium population may be smaller or larger than if there was a single driver with the ESS rate of drive, with the smaller sizes tending to occur when the stronger driver is closer to the ESS than the weaker one, and therefore more frequent.

In terms of stability, it has previously been shown in the context of a single driver and wild type that in the case of very strong drivers the equilibrium point may not be stable, and instead there is a Neimark-Sacker bifurcation (Kuznetsov, 2004) and the chromosome frequencies and population size tends to a limit cycle (Beaghton & Burt, 2022). The same dichotomy is observed here, where pairs of drivers that are very different in their rates of drive can lead to an unstable equilibrium, and pairs that are more similar tend to a stable equilibrium mixture (above and below the red line in Figs 4a,b, respectively).

The demographic parameters *R* and *α* may be difficult to measure, and in any case are likely to vary across a landscape. It is therefore interesting to see what happens if the two drivers are fixed, and *R* and *α* allowed to vary. As would be expected, in populations where *R* is higher or *α* is lower, and therefore the pre-release population density higher, the frequency of the stronger driver is higher (Fig. 5a). As a result, while the post-release equilibrium population density still increases with *R* and decreases with *α* (Fig. 5b), the extent of population suppression (relative to the pre-release equilibrium) also increases (Fig. 5c).

**Figure 5:**
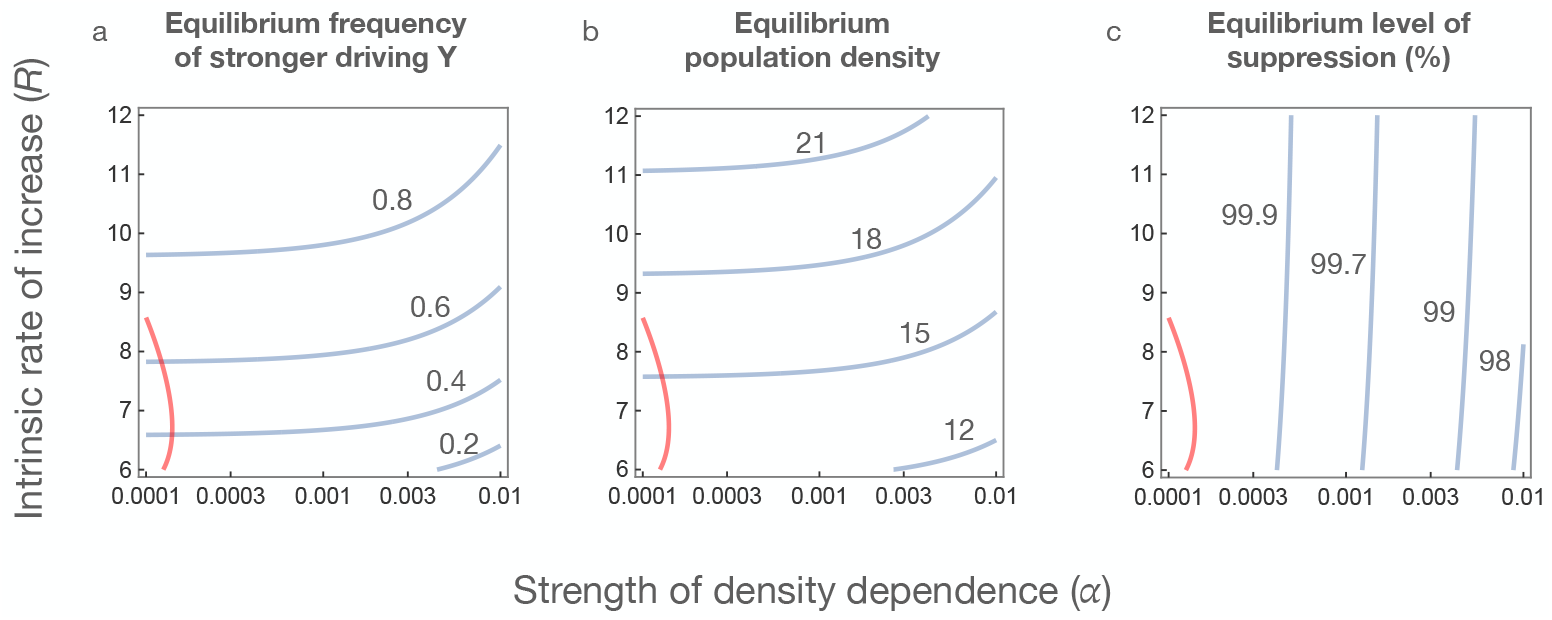
The effect of varying demographic parameters when there are two drivers straddling the ESS. Calculations are for drivers with transmission rates of *m*_*A*_ = 0.9 and *m*_*B*_ = 0.96. Contour plots show (a) the equilibrium frequency of the stronger driver, (b) the equilibrium population density, and (c) the equilibrium level of suppression (i.e., compared to the pre-release equilibrium population size) as a function of the intrinsic rate of increase (*R*) and the strength of density dependence (*α*). The red line in each plot shows the Neimark-Sacker bifurcation points separating regions where the population tends to a stable equilibrium (right of the curve) vs. a limit cycle (left).

## Inbreeding depression

In deriving Eqs(1) and (2), we assumed that all of the females have the same expected fecundity, irrespective of whether they are mated by sibling males or not. We now follow Beaghton & Burt (2022) and allow for some or all of the females mated by sibling males to be sterile and thus removed from the pool of mated females that disperse to new patches for local reproduction. We define the inbreeding depression coefficient 0 ≤ *δ* ≤ 1 as the probability of a sibling-mated female being sterile.

The generalised Eq(2) now becomes

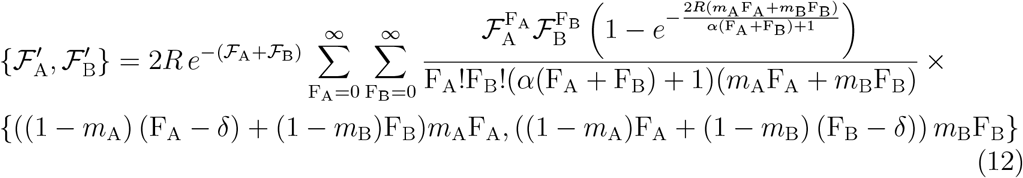

Following the same analysis as above, we derive a set of equations that can be solved numerically for *m* = *m*^∗^ and the corresponding equilibrium population *S* = *S*^∗^ for a given value of the inputs *R, α* and, additionally now, *δ*.

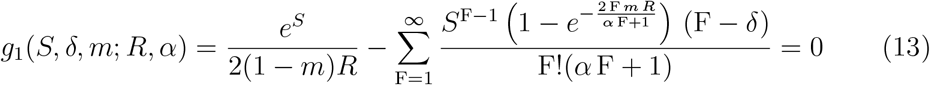

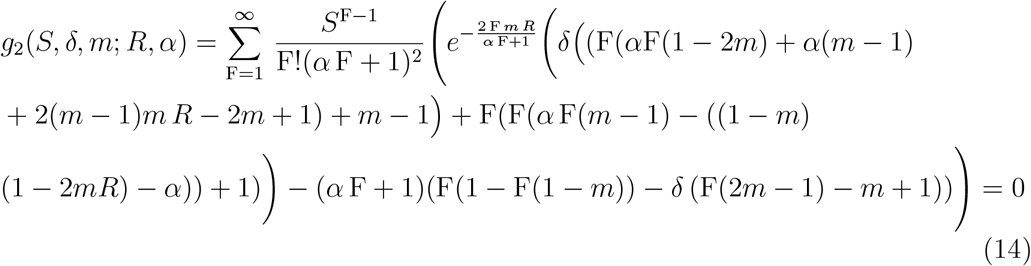

Eqs(13) and (14) reduce to Eqs(8) and (7), respectively, in the absence of inbreeding depression (i.e. when *δ* = 0).

We follow the same analysis as above to study the linear stability of the equilibrium solution 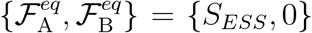, where A and B are genotypes with *m*_A_ = *m*^∗^ and *m*_B_ = *m*^∗^ + *ϵ*, respectively. Eqs(9-10) are now generalised for 0 ≤ *δ* ≤ 1 to

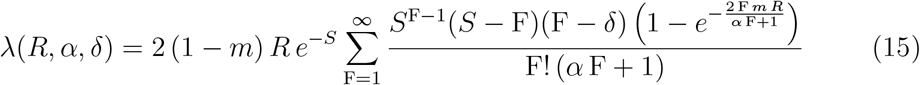

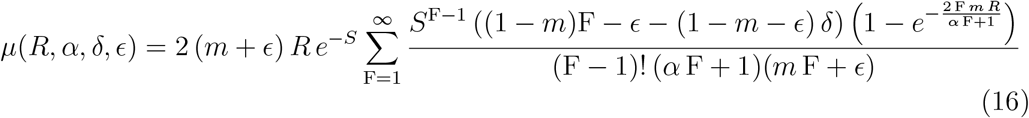

with both *m* = *m*^∗^ and *S* = *S*^∗^ also being functions of *R, α* and *δ*. As before, when |*λ*| *<* 1(*>* 1), the fixed point is linearly stable (unstable).

Note that, for given *R* and *α*, above a critical strength of inbreeding depression there is no {*m, S*} pair that satisfies Eqs(13-14), regardless of stability. We calculate this critical strength of inbreeding depression (and the associated {*m, S*} pair) by treating *S* and *δ* in Eqs(13-14) as dependent variables and *m* as an independent variable; we then implicitly differentiate Eqs(13-14) w.r.t. *S* and *m* and set 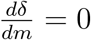:

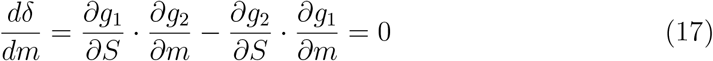

Eqs(13-14) and Eq(17) are used, for given *R* and *α*, to derive the critical *δ* (and the associated {*m, S*} pair) above which no solutions of Eqs(13-14) for *m* = *m*^∗^ and *S* = *S*^∗^ exist. For *R* = 6, *α* = 0.01, the critical value is *δ* ≈ 0.931814.

Numerical analysis of Eqs(13-14) shows that increasing inbreeding depression reduces the equilibrium sex ratio and population size, and increases instability via the eigenvalue *λ* (Fig. 6). The most important effect of including inbreeding depression is to increase the range of parameter space for *R* and *α* in which the equilibrium (*m*^∗^, *S*^∗^) pair is unstable and the population expected to evolve ever higher drive until it goes extinct (Fig. 7).

**Figure 6:**
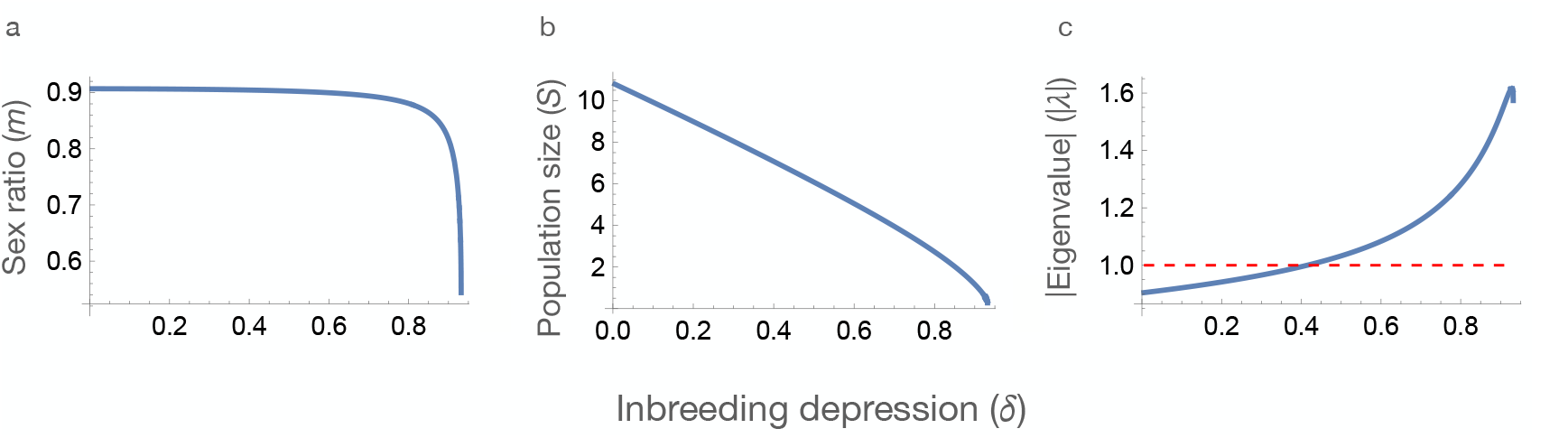
Equilibrium sex ratio, population size and magnitude of the eigenvalue (| *λ* |) as a function of the level of inbreeding depression. When | *λ* | *>* 1 (i.e., above the red dashed line) the equilibrium is unstable. *R* = 6, *α* = 0.01.

**Figure 7:**
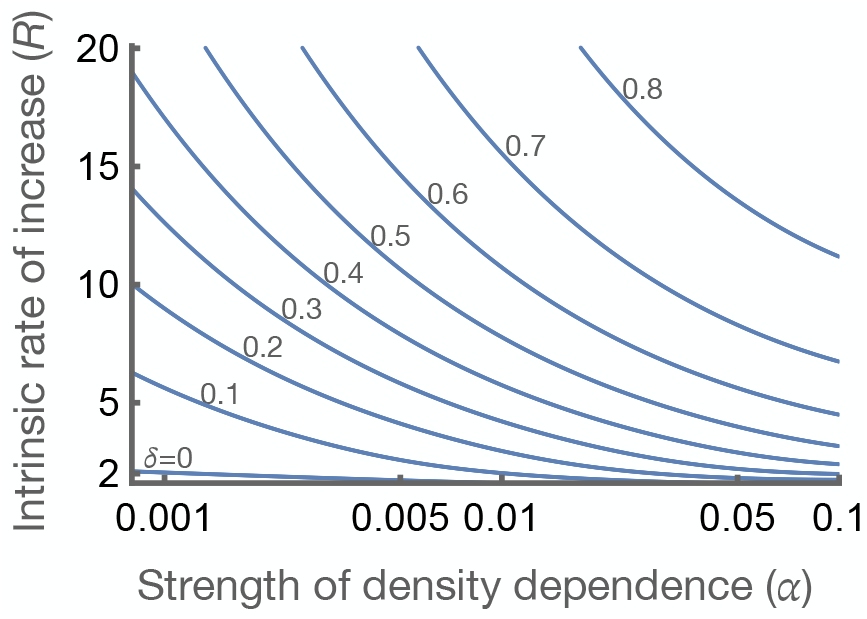
Stable vs unstable regions for different values of *δ*. Increasing the strength of inbreeding depression (*δ*) increases the size of the unstable region (i.e., the region under the corresponding line) where selection is for ever increasing drive until the population is eliminated. The line for *δ* = 0.9 is outside the area plotted. Note that the *δ* = 0 line is the red line in Fig 2.

## Discussion

The potential selective consequences of releasing a synthetic gene drive into a population has been been relatively well studied in the context of the possible evolution of resistance and how that may be retarded or prevented (Beaghton *et al*., 2017; Marshall *et al*., 2017; Unckless *et al*., 2017; Edgington *et al*., 2020; Gomulkiewicz *et al*., 2021; Cook *et al*., 2022; Khatri & Burt, 2022). The question of how selection will act on the driver itself appears to have received less attention. Variation for selection to act upon could arise by mutation and/or because multiple variants have been purposefully released.

In this paper we have studied the competitive dynamics of Y chromosomes that differ only in their rate of drive by extending an explicitly solvable spatially implicit stochastic model previously analysed by Beaghton & Burt (2022). Within the context of this modelling framework, some life histories allow the fixation of arbitrarily strong drivers, sufficient to eliminate a population, and in these scenarios we would expect evolution to always favour stronger drivers, until the population is eliminated. However, for one life history scenario, in which dispersal is by mated females, and with little or no inbreeding depression, the population persists no matter how strong the drive. This is perhaps the simplest fully stochastic model that can give indefinite persistence for all strengths of drive.

Our analysis of the extended model has shown that, in the absence of inbreeding depression, (1) for most of the biologically realistic parameter space the rate of drive will evolve to a single ESS value, and the population is substantially suppressed but persists indefinitely; (2) there is a relatively small region of parameter space (low *R* and low *α*) in which selection is for ever higher drive, until the population is eliminated; and (3) in the stable zone, if two different Y drives that straddle the ESS are released, they will typically evolve to a stable polymorphism in which, again, the population is substantially suppressed but persists indefinitely. Finally, we have shown (4) that with increasing inbreeding depression the region of parameter space in which selection favours ever stronger drive (until the population is eliminated) expands to cover most or all of the biologically realistic values.

The specific life history scenario modelled is the same as in Hamilton’s (1967) well studied local mate competition model. The key structural difference between the models is that population size (*S*) in his model is extrinsically determined, whereas in our model it is an intrinsically determined by the demographic parameters (*R* and *α*) and changes in response to the spread of a driving Y (and, in so doing, alters selection on that Y). Other differences between the two models include that he assumed each patch receives an equal number of mated females and each female produces an equal number of progeny, whereas we assume that patches receive a Poisson-distributed number of females, and each female produces a Poisson-distributed number of progeny. These latter differences appear to largely cancel out to produce approximately the same relationship between equilibrium sex ratio and population size (in the absence of inbreeding depression). The local mate competition model has been widely used to study the evolution of the sex ratio when under maternal control, such as in haplo-diploid species (reviewed in West, 2010), and these models could also be expanded to consider the joint evolution of sex ratio and population size, with stability assessed.

Previous work demonstrated that, in the absence of inbreeding depression, if a driving Y is released that is “too strong” for the given demographic parameters, then the population size does not tend to a single stable value, but instead to a limit cycle, which may not be desirable in the context of population management (Beaghton & Burt, 2022). The problem is exacerbated because the key demographic parameters are difficult to measure, and are expected to vary over space and time, making it difficult to release a driver with the “right” strength of drive. Any driver that is released would be expected to evolve over time by the selection of *de novo* mutations to the ESS rate of drive, which would stabilise population size, but this could take a long period of time, depending on the mutation rate. In principle, this process could be accelerated by incorporating a hyper-mutable region in the construct that affects the rate of drive (e.g., a di- or tri-nucleotide spacer between promoter and coding sequence; Simoni *et al*., 2020). Here we have explored a potentially simpler alternative, in which two driving Y’s are released that straddle the ESS. Their frequencies in the population can relatively rapidly go to equilibrium values which produce an overall sex ratio and level of population suppression that is not too different from the ESS. Moreover, these frequencies can be expected to adjust over space and time to changes in the underlying demographic parameters, and in this way the likelihood of population size cycles reduced.

Increasing inbreeding depression increases the parameter space in which ever stronger drive is selected for, until the population is eliminated, because it nullifies the low-density advantage that lower drive males have more sisters to mate with (Beaghton & Burt, 2022). Inbreeding depression therefore has a selective effect on the proportions of the two drivers in the population. Inbreeding depression also acts as an Allee effect, reducing population growth rates at low density, but the relative importance of these two factors (selection and overall growth rates) is unclear and it would be interesting to investigate the stabilising/destabilising influence of other factors that reduce population growth rates at low density but do not have the same selective effect as inbreeding depression. Most potential targets of control by gene drive are outcrossing (in part because gene drive does not work efficiently in highly inbred species), and can therefore be expected to show considerable levels of inbreeding depression.

Other potential extensions of this work include: (1) examining the dynamics of alternative drivers in spatially explicit and spatially realistic models (i.e., in which some pairs of populations are closer together (and share more migrants) than others; Eckhoff *et al*., 2017; North *et al*., 2020; Birand *et al*., 2022; Liu *et al*., 2023; Zhu & Champer, 2023); (2) extending to other types of drive, such as autosomal homing or toxin-antidote drivers, which typically differ from Y drives in not going to fixation even in non-spatial models; and (3) investigating the interactions of drivers at different loci (e.g., a Y-linked driver and an autosomal driver, or two autosomal drivers at different loci). Given the possibilities of either multiple drivers being released into a species, or a single driver mutating to form alternative variants, it is important to have a good theoretical understanding of the likely consequences.

## Appendices

### Appendix A: Arbitrary number of driving Ys

Here we show the recurrence equation for an arbitrary number of *N ≥* 2 male types, each with a different strength of drive 0 ≤*m*_*i*_ *≤*1, 1 ≤*i ≤N*. Following the same mathematical approach, we derive the recurrence equations for the evolution over one generation of the population densities **ℱ**= {ℱ_1_, *ℱ*_2_, …, *ℱ*_*N*_}of the *N* mated female types in the cloud, given the strengths of drive {*m*_1_, *m*_2_, …, *m*_*N*_}and the inputs *R* and *α*:

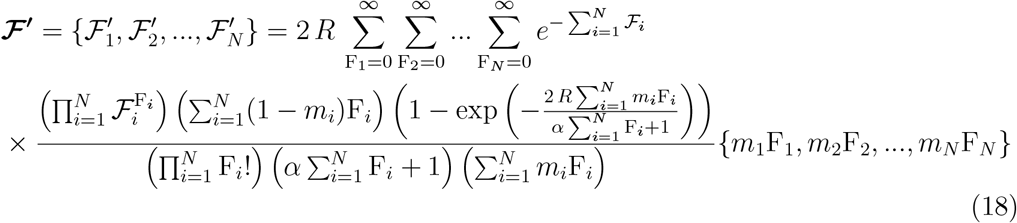

We postulate that of the *N* male types introduced, as *t*→ ∞, **at most** two types will survive, i.e. either one will tend to fixation or two will coexist.

### Appendix B: Single genotype system

Here we analyze, using Eq(12), a single genotype system with strength of drive *m*_A_ = *m* (we simply set ℱ_B_ = 0). The equation for the equilibrium population density 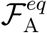 (denoted here as *S*) is

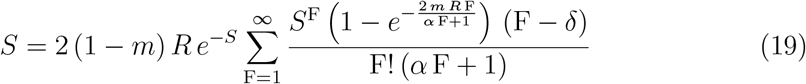

Note that this equation reduces to Eq(8) when *δ* = 0. In addition to the trivial solution *S* = 0, for given {*R, α, m, δ*}, Eq(19) admits one or two solutions with *S >* 0. We now examine the linear stability of these equilibrium solutions:

#### B1: The trivial fixed point *S* = 0

We differentiate the RHS of Eq(19) wrt *S* and set *S* = 0. For values of {*R, α, m, δ* = 0} where this (always positive) derivative, the eigenvalue 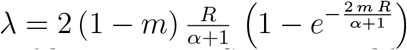 of the linearised equation, is less than 1, the trivial equilibrium state *S* = 0 is stable and thus the introduction of a small population density in an empty landscape fails. Conversely, when *λ >* 1, the state is unstable and the population grows after the initial introduction. Setting *λ*(*R, α, m, δ* = 0) = 1 allows us to calculate for any given 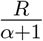 the critical *m* above (or below) which the trivial state is stable (or unstable). The solution of equation 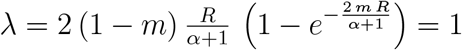 is the critical 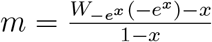, where 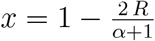 and *W*_*r*_ (*y*) is the *r*-Lambert function. The analysis above is qualitatively the same for *δ >* 0.

#### B2: Fixed point(s) *S >* 0

We can only solve Eq(19) numerically for given {*R, α, m, δ*} (there can be one or more solutions *S >* 0) and we then evaluate the derivative

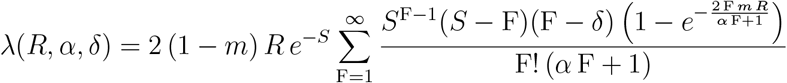

of the RHS of Eq(19) at *S >* 0. The equilibrium state is again linearly stable when | *λ*| *<* 1. Note that the above eigenvalue for the single genotype system is one of the two eigenvalues of the two genotype system, given by Eq(15).

#### B3: The unstable region for *S >* 0

In the unstable region in Fig. 2 (and the corresponding figures for non-zero values of *δ*), for any given value of *R, α*, and *δ*, and depending on the strength of drive *m*, there are zero, one, or two non-zero solutions for the equilibrium population *S >* 0 of the single-genotype system.

We rearrange Eq (19):

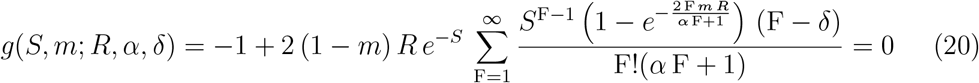

Eq(20) can be solved for *S*, given *m* and the parameters *R, α*, and *δ*. The expression *g*(*S, m*; *R, α, δ*) in Eq(20) is a concave function of *S* and has a single root wrt *S* when 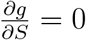:

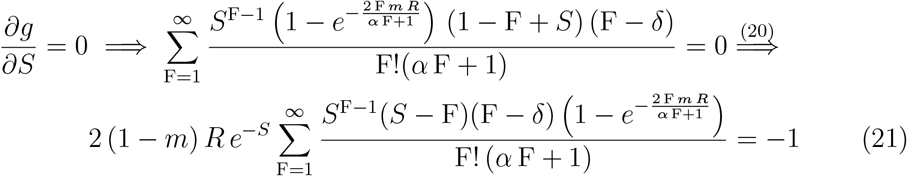

Eqs (20) and (21) yield *m* and *S* for given *R, α*, and *δ*. For all values of drive less than *m* there are two non-zero equilibrium values of *S* whereas for values of drive greater than *m* there are no non-zero equilibrium values of *S*.

### Appendix C: Selection gradient in the stable region: change of sign at ESS

In Eqs(1) we set {*m*_*A*_, *m*_*B*_}= {*m* + *ϵ/*2, *m − ϵ/*2}, {ℱ_A_, *ℱ*_B_}= {*S p, S* (1 − *p*)}, where *S* is the stable equilibrium population size associated with the value of *m* (from Eq 8) and *p* is the frequency of the stronger driver. We now have the following equations for 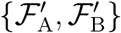 in the next generation:

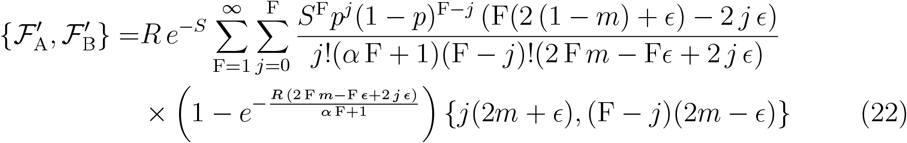

The frequency of the stronger driver (A) in the next generation is defined as 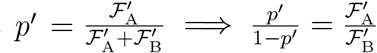. From Eq(11), the selection gradient becomes

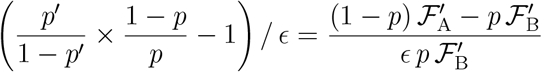

Eqs(22), along with the definition of the selection gradient above are used to generate the plots in Fig 3. Further, to identify the value of *m* where the selection gradient changes sign, we set the selection gradient equal to zero, which results in

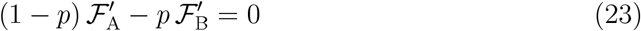

We now combine Eq(22) and Eq(23):

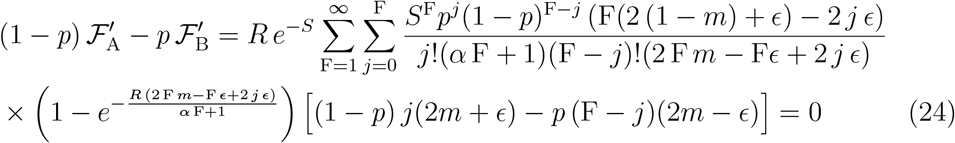

We focus on small values of *ϵ* and expand the double summation term in Eq(24) as an infinite series in *ϵ* about *ϵ* = 0. The zeroth order term turns out to be identically zero. The first order term (i.e. the one proportional to *ϵ*) reduces to an single infinite sum

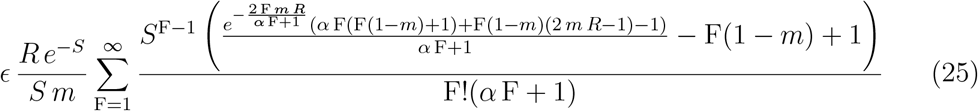

As *ϵ* → 0, we can disregard higher order terms in the series and from Eqs(24-25) we obtain

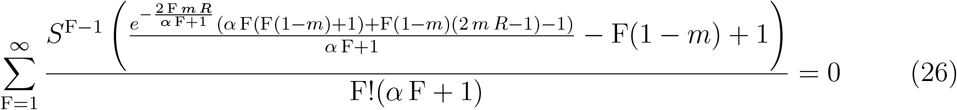

We had chosen S to be the stable equilibrium population size associated with the value of *m* so it satisfies Eq(8):

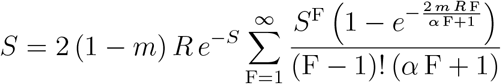

Eq (26) is identical to Eq(7) so when solved in conjunction with Eq(8), it gives a unique pair of ESS values {*m* = *m*^∗^, *S* = *S*^∗^} in the stable region of *R* and *α*.

We have thus shown that the critical value of *m* where the selection gradient changes sign in the stable region is the one that corresponds to the ESS.

## Acknowledgements

This work was supported by the Bill & Melinda Gates Foundation and the Open Philanthropy Project.

